# Epidemiology of the Zika virus outbreak in the Cabo Verde Islands, West Africa

**DOI:** 10.1101/198952

**Authors:** José Lourenço, Maria de Lourdes Monteiro, Tomás Valdez, Júlio Monteiro Rodrigues, Oliver G. Pybus, Nuno Rodrigues Faria

## Abstract

**Introduction:** The Zika virus (ZIKV) outbreak in the island nation of Cabo Verde was of unprecedented magnitude in Africa and the first to be associated with microcephaly in the continent.

**Methods:** Using a simple mathematical framework we present a first epidemiological assessment of attack and observation rates from 7,580 ZIKV notified cases and 18 microcephaly reports between July 2015 and May 2016.

**Results:** In line with observations from the Americas and elsewhere, the single-wave Cabo Verdean ZIKV epidemic was characterized by a basic reproductive number of 1.85 (95% CI, 1.5 −2.2), with overall the attack rate of 51.1% (range 42.1 - 61.1) and observation rate of 2.7% (range 2.29 - 3.33).

**Conclusion:** Current herd-immunity may not be sufficient to prevent future small-to-medium epidemics in Cabo Verde. Together with a small observation rate, these results highlight the need for rapid and integrated epidemiological, molecular and genomic surveillance to tackle forthcoming outbreaks of ZIKV and other arboviruses.

## Main Text

Zika virus (ZIKV) was first reported in Uganda in 1947 but until recently human infections in Africa were considered rare and not associated with outbreaks or neurological complications (1). The first large ZIKV outbreak was reported in the Yap island of Micronesia in 2007 where around 73% of the population is estimated to have been infected (2). In 2013-14, a similar proportion of the population was exposed to the virus in French Polynesia, where the first ZIKV-infections associated with neurological complications were detected (3). In early May 2015, autochthonous transmission of the virus was confirmed in the northeast region of Brazil (4). Subsequent analysis of genetic data suggests that ZIKV had been circulating undetected in the Americas at least since early 2014 (5, 6). In February 2016, ZIKV became a Public Health Emergency of International concern, and two months later the World Health Organization (WHO) and the Centre for Disease Control (CDC) confirmed the association of Zika virus infection with microcephaly and other neonatal neurological complications. So far, of the two known Zika virus lineages – the African genotype, restricted to the African continent, and the Asian genotype, confined to Southeast Asia and the Americas – only the later has been associated with microcephaly and neonatal neurological complications.

Recently, the archipelago of Cabo Verde reported 7,580 ZIKV suspected cases and 18 microcephaly cases between October 2015 and May 2016, in the first known ZIKV outbreak in Africa. Cabo Verde is composed by 10 islands located west of the coast of Senegal, with a total population size of 539,560 in 2016. Importantly, this is also the first time that ZIKV-associated microcephaly cases are reported in Africa. While epidemiological investigations based on clinical reports in the Americas may have been obscured by the co-circulation of dengue and chikungunya viruses – which can confound case report data due to overlapping clinical symptoms with ZIKV; these were not reported to be co-circulating in Cabo Verde at the time of the ZIKV outbreak. Despite its scientific and public health importance, there is a dearth of information about the Cabo Verdean ZIKV outbreak. Here we provide the first epidemiological characterization of the outbreak in Cabo Verde in light of the information gathered from the recent epidemics in Brazil and French Polynesia.

By the 5th of October 2015 (epidemiological week 40), the first cases of illness with skin rash were reported in the capital city of Praia, on the island of Santiago. A detailed report is available (in Portuguese) from the official Surveillance and Outbreak Response Unit of the Ministry of Health of Cabo Verde (SVIER-MS) (7). ZIKV suspected case counts were based on clinical evaluation following Cabo Verde’s definition of patients presenting exanthema (skin rash) with or without fever and at least one of the following symptoms: arthralgia (joint pain), myalgia (muscle pain) or conjunctivitis (not purulent and without hyperemia) (8). The number of ZIKV cases grew rapidly, and by the end of week 41 (2015) there were 95 ZIKV notified infections (Figure 1a). The peak of the outbreak occurred in week 47 (22-28th November, 2015). The outbreak was characterized by a single epidemic wave and transmission ceased in week 21 of 2016 (22-28th May).

**Figure 1.**
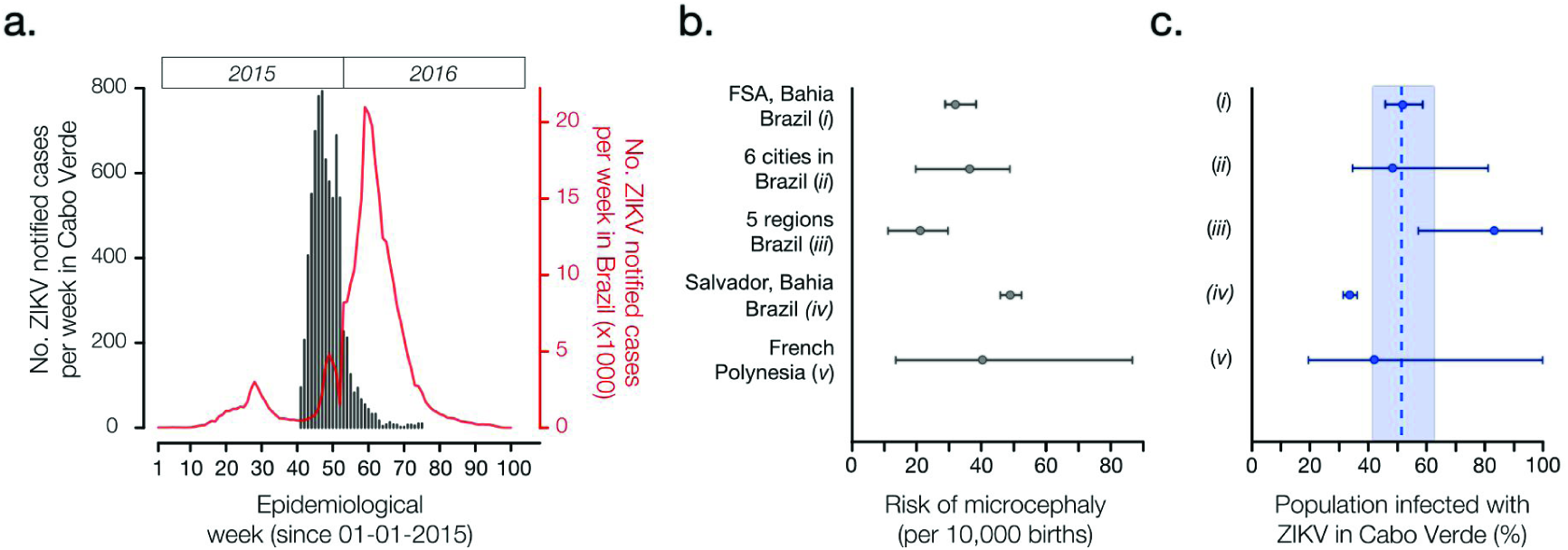
Epidemiological characteristics of Zika virus outbreak in Cabo Verde during 2015-2016. (Panel a) Number of ZIKV notified cases per epidemiological week in Cabo Verde (dark grey) and in Brazil (red). (Panel b) Reported number of microcephaly cases per 10,000 ZIKV-exposed pregnancies and corresponding 95% confidence intervals by studies (i)-(v). (Panel c) Estimated proportion of the Cabo Verdean population infected with ZIKV during the 2015-2016 outbreak (attack rate, AR), as formulated by AR = m /(b x rMC). We considered m=18 microcephaly cases across Cabo Verde as reported to the SVIER-MS, and b=10,908 births during the observation period; rMC=risk of microcephaly per pregnancy extrapolated from reports in panel b. Dashed line and coloured area represent the mean and standard error estimates for the AR in Cabo Verde.

By 29th of May 2016, 7,580 suspected ZIKV cases had been reported by health centers in four out of ten islands (Santiago, Fogo, Maio, Boavista). Two of these islands reported most cases: Santiago reported 65.1% (4937/7580) and Fogo 19.2% (1458/7580). A total of 18 microcephaly cases were confirmed; all microcephaly cases occurred within the four islands with reported ZIKV transmission (12 in Santiago, 4 in Fogo, 2 in Maio). Microcephaly cases were confirmed based on ecography and/or measured cephalic perimeter of &#x3C;32 cm of the newborn (detailed in 9). Approximately 50% of all confirmed microcephaly cases were linked to reports of Zika-related symptoms in the first trimester of gestation. Sixty-four samples related to suspected Zika cases were sent to Pasteur Institute in Dakar; of these, 17 tested positive for Zika (15 were IgM positive, 2 were RT-qPCR positive); all samples tested negative for dengue, chikungunya, Yellow fever, Rift Valley fever and West Nile (10).

We first estimated the basic reproductive number (R0) for the Cabo Verde outbreak, defined as the average number of secondary human cases generated by a single primary human case in a totally susceptible population. Based on weekly case counts reported by the SVIER-MS (Figure 1a) (7), we fitted a simple exponential growth model to the weekly number of suspected ZIKV cases (weeks 40-46), using a gamma distribution for generation times, as previously described and implemented for Brazil regions (5). The epidemic generation time was assumed to have mean 10.8 and standard deviation of 3.9 days, as recently estimated for Feira de Santana (Brazil) (11). Our model revealed an R0 with mean 1.85 (95% CI, 1.5 - 2.2) for the Cabo Verde outbreak. This was in line with R0 values obtained for different regions in Brazil (varying from 1.29 to 1.98 (5)).

Next, we sought to quantify the attack rate (AR) of ZIKV in Cabo Verde, defined as the proportion of the entire population infected with the virus during the first epidemic wave. We calculated the AR based on the observed number of microcephaly cases (m), observed number of newborns in Cabo Verde (b), and estimated absolute risk of microcephaly per pregnancy (rMC). Here, the expression for the attack rate is simply AR=m/(b x rMC). We considered m=18, the number of reported microcephaly cases in Cabo Verde as reported to the SVIRE-MoH, and b=10,908 live births during the same period (1 year).

We calculated the AR in Cabo Verde under five different absolute risks of microcephaly per full pregnancy (rMC) from studies based in regions of Brazil and the French Polynesia (Figure 1b, 1c). Study (i) used a climate-driven transmission model to investigate the ZIKV-related absolute risk of microcephaly in Feira de Santana (FSA), the second largest municipality in Bahia state, Northeast Brazil; assuming an AR=65%, the number of of microcephaly cases in FSA was estimated at 31.9 (95% CI: 28 to 36) cases per 10,000 challenged pregnancies (11); this resulted in an extrapolated AR= 51% (95% CI: 45 to 58) for Cabo Verde. Study (ii) analysed data from 6 municipalities in 3 states of Brazil, and estimated 34.2 (95% CI: 20 to 48) microcephaly cases per 10,000 challenged pregnancies, with an assumed AR=50% (12); in this case, the extrapolated AR for Cabo Verde was 48% (95% CI: 34 to 81). Using official numbers from the Brazilian Ministry of Health, study (iii) reported 19.8 (95% CI: 10 to 29) microcephaly cases per 10,000 challenged pregnancies (13), which returned a ZIKV AR of 83% (95% CI: 56 to 100) for Cabo Verde. Study (iv) estimated a ZIKV AR of 69.3% on pregnancies not deriving into neonate neurological pathologies and reported 49.08 (95% CI: 46.5 to 52.3) microcephaly cases per 10,000 challenged pregnancies in Salvador, Bahia during the 2015-2016 epidemic (14); using this information, we estimated an AR for Cabo Verde of 33.6% (95% CI 31.5 to 35.4). Finally, study (v) estimated 40 (95% CI: 13 to 86) microcephaly cases per 10,000 challenged pregnancies during the 2013-2014 outbreak in the French Polynesia, characterised by an AR=66% (95% CI: 62 to 70) (3); this resulted in an extrapolated AR=41% (95% CI: 19 to 100) for Cabo Verde.

Overall, the mean attack rate of ZIKV in Cabo Verde was estimated at 51.1% (standard error, SE, 42.1 - 61.1) when considering the independent estimations from studies (i) to (v). This mean (Figure 1c) is slightly lower than for previous outbreaks (2, 3, 11, 12, 14), suggesting that the level of herd-immunity may still be under the threshold that would prevent additional small to medium size ZIKV outbreaks in Cabo Verde. Given the estimated AR and the number of reported infections, we estimate a low case observation rate of OR=2.7% (SE 2.29 to 3.33), i.e. ~27 cases were notified for every 1,000 infections. A low OR is not uncommon for ZIKV (see for instance Table 3 of (11)).

With an AR ranging from 42% to 61%, our findings suggest that between ~221,000 to ~329,000 people may have been infected with ZIKV during the recent outbreak in Cabo Verde. Was the viral lineage that caused this large outbreak in Cabo Verde introduced from the Americas or from another African country? A rapid assessment of the global air travel network suggests the presence of similar ecologies and direct air travel connections between the northeast of Brazil and Santiago island where most cases were observed. Given the timing of the epidemic (Figure 1a) and the high number of travellers visiting Cabo Verde from the Americas it seems plausible to speculate that the outbreak was caused by the Asian genotype circulating in the Americas (the country received >7000 travellers from ZIKV infected countries in 2015, including direct flights from Northeast Brazil (15)). Moreover, recent ZIKV cases in Africa have also been reported in Angola (1 returning traveller to France and 1 autochthonous case (16)) and Guinea-Bissau (3 cases in the Bjagó islands (17)). Similar to Cabo Verde, no genetic data is available from Angola or Guinea-Bissau.

Our mathematical framework is based on two fundamental assumptions: (i) that the AR among pregnant women is the same as the in general population, and (ii) the risk of MC without prior exposition to ZIKV is negligible. The effect of these assumptions in our AR estimations for Cabo Verde are expected to vary between data sources. Without the support of more local data these assumptions and their effects are however difficult (if not impossible) to assess. Reassuringly, we note that the independent estimation of R0 using the exponential phase of the epidemic predicts an AR=1-1/R0 of mean 46% (95% CI 33 - 54) with significant overlap with the estimation obtained from assumptions (i) and (ii) (51.1%, SE 42.1 - 61.1).

Recent advances in portable genome sequencing allow to generate genome data in the field within days (18, 19), and can help increasing research capacity, thus reducing time to response to future outbreaks. In the future, retrospective seroepidemiological surveys could further facilitate the estimation of attack, symptomatic and observation rates not only for ZIKV but also for other arboviruses. In 2009, Cabo Verde reported >17,000 suspected infections with dengue virus (20) in a single epidemic similar to others (21). Thus, Cabo Verde’s epidemiological setting is unique given that dengue and Zika have caused single but sequential epidemic waves there. This offers an exceptional opportunity to evaluate the potential contribution of previous dengue virus seropositivity to the ZIKV associated risk of microcephaly.

In conclusion, our early epidemiological assessment of the largest ZIKV outbreak in Africa suggests that half of the Cabo Verde population was exposed in 2015-2016. Since other regions have consistently reported higher attack rates, it is possible that Cabo Verde could still witness a second ZIKV outbreak. This finding highlights the need for an integrated epidemiological, molecular and genomic arbovirus surveillance system in order to improve control in forthcoming outbreaks in Cabo Verde.

